# GTRpmix: A linked general-time reversible model for profile mixture models

**DOI:** 10.1101/2024.03.29.587376

**Authors:** Hector Banos, Thomas KF. Wong, Justin Daneau, Edward Susko, Bui Quang Minh, Robert Lanfear, Matthew W. Brown, Laura Eme, Andrew J. Roger

## Abstract

Profile mixture models capture distinct biochemical constraints on the amino acid substitution process at different sites in proteins. These models feature a mixture of time-reversible models with a common set of amino acid exchange rates (a matrix of exchangeabilities) and distinct sets of equilibrium amino acid frequencies known as profiles. Combining the exchangeability matrix with each profile generates the matrix of instantaneous rates of amino acid exchange for that profile.

Currently, empirically estimated exchangeability matrices (e.g., the LG or WAG matrices) are widely used for phylogenetic inference under profile mixture models. However, such matrices were originally estimated using site homogeneous models with a single set of equilibrium amino acid frequencies; therefore unlikely to be optimal for site heterogeneous profile mixture models. Here we describe the GTRpmix model, implemented in IQ-TREE2, that allows maximum likelihood estimation of a common set of exchangeabilities for all site classes under any profile mixture model. We show that exchangeability matrices estimated in the presence of a site-heterogeneous profile mixture model differ markedly from the widely used LG matrix and dramatically improve model fit and topological estimation accuracy for empirical test cases.

Because the GTRpmix model is computationally expensive, we provide two exchangeability matrices estimated from large concatenated phylogenomic supermatrices under the C60 profile mixture model that can be used as fixed matrices for phylogenetic analyses. One of these, called Eukaryotic Linked Mixture (ELM), is designed for phylogenetic analysis of proteins encoded by nuclear genomes of eukaryotes, and the other, Eukaryotic and Archeal Linked mixture (EAL), for reconstructing relationships between eukaryotes and Archaea. These matrices when combined with profile mixture models fit data much better and have improved topology estimation relative to the empirical LG matrix combined with the same underlying mixture models. Version v2.3.1 of IQ-TREE2 implementing these models is available at www.iqtree.org.

## 1 Introduction

Models of amino acid substitution are of key importance to probabilistic molecular phylogenetic analyses of protein sequences. Typically, the amino acid substitution process is modeled via a site-independent time-reversible Markov process on a tree. The parameters of this model include a set of fixed equilibrium frequencies of the amino acids (also known and referred to in this work as a profile) and a constant matrix of amino acid exchange rates – exchangeabilities – throughout the tree. The exchangeability matrix accounts for some biological, chemical, and physical amino acid properties and when combined with the equilibrium frequencies of the amino acids, it describes the instantaneous rates of interchange between each pair of amino acids.

In most phylogenetic analyses the exchangeability matrix used is chosen from a set of fixed empirically-estimated matrices. The first empirically estimated exchangeabilities were derived from the Dayhoff (Dayhoff et al., 1978) and Jones-Taylor-Thornton (JTT) (Jones et al., 1992) matrices that were obtained by counting the substitutions between each amino acid pair using ancestral sequence reconstruction and a parsimony-based approach to analyze databases of multiple alignments along with their estimated phylogenies. Subsequently, a maximum likelihood approach was used to improve exchangeability estimation leading to the development of the “Whelan and Goldman” (WAG) model (Whelan and Goldman, 2001). Le and Gascuel expanded this approach (Le and Gascuel, 2008) by considering larger data sets and by incorporating heterogeneity of rates across sites in the likelihood computation via a site-rate partition model. The resulting matrix, known as the ‘Le and Gascuel’ (LG) matrix, is currently very widely used for phylogenetic inference based on protein sequences. Expanding on these, Minh et al. introduced QMaker (Minh et al., 2021), a maximum likelihood method to estimate an exchangeability matrix from a large protein data set consisting of multiple independent sequence alignments. The authors used QMaker to estimate a number of additional matrices to be used for phylogenetic analyses of specific taxonomic groups (e.g. Q.bird, Q.insect, and Q.plant). Other matrices have been developed to fit proteins encoded on certain organellar genomes (e.g. cpREV (Adachi et al., 2000)) or particular gene families (e.g. rtREV (Dimmic et al., 2002)).

All of the foregoing exchangeability matrices were obtained assuming that all sites evolve according to the same process and share a single set of equilibrium amino acid frequencies (a single profile). However, because of different functional constraints and structural microenvironments within proteins, there are distinct ranges of admissible amino acids at sites (Franzosa and Xia, 2009; Goldstein, 2008; Pál et al., 2006). Profile mixture models, such as the C10-C60 series and the UDM series (Schrempf et al., 2020; Si Quang et al., 2008; Wang et al., 2008, 2014), were designed to account for this heterogeneity of preferred amino acids across sites. These models are mixtures of time-reversible Markov models, but they assume a common exchangeability matrix and distinct profiles of stationary frequencies.

Estimation of a single exchangeability matrix, within a profile mixture setting, has been explored in a Bayesian context by Lartillot and colleagues (Lartillot and Philippe, 2004) through the development of various versions of the CAT model of Phylobayes (Lartillot et al., 2013). In the maximum likelihood framework, Wong and colleagues (2022) developed MAST (Wong et al., 2024), an extension of IQ-TREE2 that, among other things, allows the user to estimate a mixture model with various options for linking and unlinking exchangeability matrices and amino acid profiles, in conjunction with mixtures of tree topologies. While this implementation can be very useful in many contexts, it is not practical for profile mixture models with many profiles because, for each profile, 189 exchangeability parameters need to be estimated. For commonly used models with 40 to 60 profiles (e.g. C40 or C60) or more (e.g. UDM64, UDM256, etc.), this corresponds to *>>*7500 estimated parameters. These models would require complex and computationally expensive optimization and will potentially be susceptible to problems associated with local optima, over-parameterization, and identifiability.

Here we describe the implementation of a General-Time-Reversible model via maximum likelihood estimation in IQ-TREE2 for use with profile mixture models. This GTR model (denoted as GTR20 in IQ-TREE2) has a single set of optimizable exchangeability parameters shared (‘linked’) over all classes of the profile mixture. By simulation, we show that our implementation accurately estimates exchangeability parameters and that it can improve tree topology estimation accuracy. Additionally, we show that the estimation of exchangeabilities under a profile mixture model provides a much-improved fit on a well-known empirical data set than the profile mixture model with LG exchangeabilities.

Since the estimation of exchangeabilities can be computationally expensive and requires large data sets for accurate parameter estimates, we provide two matrices estimated from large concatenated supermatrices under the GTR-C60 profile mixture model to be used as fixed matrices for phylogenetic analyses. One of these, called ELM, is tailored for phylogenetic analyses of proteins encoded by eukaryotic nuclear genes, and the other, EAL, is for reconstructing relationships between eukaryotes and Archaea. We show, via three well-known empirical data sets, that these matrices have better fit and topological accuracy than the LG matrix when both are combined with C60. Additionally, we show that these matrices perform well with different sets of profiles.

## 2 Profile Mixture Models and Exchangeability Optimization

The general time-reversible model (GTR) (Tavaré, 1986) is a Markov process where, for a profile ***π*** = (*π*_1_, *π*_2_, …, *π*_20_), with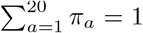, and a matrix *Q* of instantaneous rates of change between amino acids, diag(*π*)*Q* = *Q*^*T*^ diag(*π*). Because of this, one can parameterize *Q* via a non-negative symmetric matrix *S* known as the exchangeability matrix. Specifically, given an exchangeability matrix 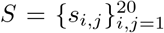 and a profile ***π***, the entries of the time-reversible instantaneous rate matrix 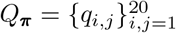 associated to ***π*** are defined as

1. *q*_*ij*_ = *s*_*ij*_*π*_*j*_, for *I j* and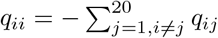, otherwise.
2. multiplying all entries by 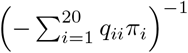, so that branch lengths are interpretable as expected number of substitutions per site.

All entries of *Q*_***π***_ are non-negative, row sums are 0, ***π****Q*_***π***_ = 0, and diag(***π***)*Q*_***π***_ is symmetric. For any *c >* 0, exchangeability matrices *S* and *cS* yield the same instantaneous rate matrix *Q*_***π***_, and thus produce the same site-pattern probabilities. Therefore we constrain one entry to be equal to 1 resulting in 189 free parameters from the exchangeability matrix.

Profile mixture models are mixtures of time-reversible models with a common exchangeability matrix *S*. Specifically, site profiles ***π***_*c*_ are selected independently with probability *w*_*c*_ and, independently of these, rates for sites, *r*_*k*_, are chosen with probability *d*_*k*_. Given the rates and site profile for a site *p*, the evolutionary model is a GTR process with exchangeability matrix *S* along a tree *T*. Let *P* (***x***_*p*_|*T, S*, ***π***_*c*_, *r*_*k*_) denote the conditional probability of site pattern ***x***_*p*_ given its site rate, *r*_*k*_, and site profile, ***π***_*c*_. Because the rates and site profiles are unobserved, the likelihood contribution under the model for the site is the marginal probability of the site pattern,

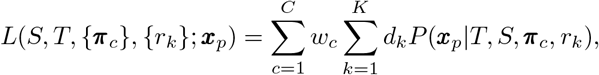

were 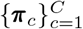 is a collection of *C* profiles with corresponding positive weights 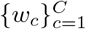 summing to one, 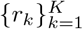 is a collection of *K* non-negative scalar rate parameters with corresponding positive rate weights 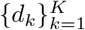 also summing to one, and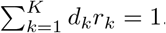.

To reduce the complexity and computational cost of profile mixture models, fixed profiles are typically used for tree estimation. In these cases, the only additional parameters coming from the profile mixture are the weights of the profile, giving *C−*1 additional free parameters from the weights, where *C* is the number of profiles. Different sets of profiles have been estimated via databases of alignments. For example, Quang and colleagues (Si Quang et al., 2008) introduced the widely used sets of profiles known as C10, C20, C30, C40, C50, and C60 (generically referred to as CXX). In each of these, the number next to the ‘C’ denotes the number of profiles in the set. In most our fits of the C-series models below we estimate the weights of the mixture; the default of IQ-TREE is to use empirical weights (weights obtained during the estimation of the original empirical profile frequencies, rather than being re-estimated during a later analysis) unless a ‘+F’ component is included. Other sets of profiles include the more recently introduced UDM models (Schrempf et al., 2020), with sets of profiles ranging from 4, 8, 16, and up to 4096 elements.

For rates across sites, it is common to use a discretized approximation to the gamma distribution with shape parameter *α* and mean 1 (Yang, 1994). For these distributions, all rates have an equal probability of occurrence and are continuous functions *r*_*k*_(*α*) of *α*. The shape parameter *α* adds only one free parameter to a profile mixture model. The discretized gamma distribution is commonly discretized into 4-rate classes and is denoted G4.

Given an MSA with *n* sites and a tree *T*, we estimate the exchangeabilities by maximizing the log-likelihood across all sites

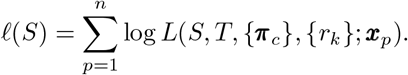

To do this, we arbitrarily fix the exchangeability between *Y* and *V* (corresponding to the entry *s*_19,20_ of *S*) to 1, and we then estimate the 189 remaining exchangeabilities using BFGS-algorithm (Fletcher, 1987), a well-known iterative optimization method. By default, the algorithm is initialized with all 189 exchangeabilities equal to one, with the option to specify any other initial exchangeabilities. In its current implementation, other parameters of the profile mixture can be jointly estimated using IQ-TREE2’s routines. For example, one can simultaneously estimate the tree topology, branch lengths, rates (not necessarily from a discretized gamma G*K*), weights of fixed profiles, and exchangeabilities (or any subset of this list).

We compare exchangeabilities *S* and *S*^*′*^ via their associated rate matrices *Q* and *Q*^*′*^ under the uniform profile; *π*_*i*_ = 1*/*20. Under this transformation *Q*_*ij*_ *∝ S*_*ij*_ is a function of *i* and *j*, so the rate matrix entries can be thought of as exchangeabilities and we refer to them as such. But the transformation to *Q* and *Q*^*′*^ puts the exchangeabilities onto a more comparable scale, one that is more closely associated with their end-use in rate matrices than setting one entry to 1 as was done in optimization.

## 3 Data Sets

Because of the computational burden associated with the estimation of the exchangeabilities in the GTRpmix model, we have analyzed two large concatenated protein ‘super-matrix’ datasets to estimate ‘general use’ substitution matrices for phylogenetic estimation with profile mixture models. These can be used with profile mixture models when sufficient computational resources are not available for full GTRpmix optimization or the datasets to be analyzed are too small to allow accurate estimation. The two datasets used to estimate these matrices are a pan-eukaryotic concatenated supermatrix and a eukaryote-archaea supermatrix.:

### Pan-Eukaryotic data sets

To estimate the “Eukaryote” exchangeability matrix we selected a 240-protein data set with 76,840 sites and 78 taxa as a taxonomically representative subsample of all eukaryotes in the PhyloFisher database (Tice et al., 2021). Taxa were selected based on their known membership in a particular higher-level eukaryotic taxon and their phylogenetic position. Further selection was done to maximize gene coverage within the original PhyloFisher data set. As detailed below, to compare two methods of exchangeability estimation, we also looked at a smaller subset of the PhyloFisher database consisting of a 240-protein data set with 77,965 sites 50 taxa.

### Eukaryotic-Archaeal data set

To estimate the exchangeability matrix for reconstructing relationships between eukaryotes and Archaea, we used a 54-protein data set with 14,704 sites and 86 taxa. This data set includes a subset of the taxa presented in Eme et al. (2023).

For more details on the datasets and their taxonomic selection see the Supplement’s section *‘Data sets*.*’*

### 3.1 Data Sets for Comparisons

The following data set is used to compare the fit of the new empirically estimated matrices to be used for reconstructing relationships between eukaryotes and Archaea against the LG matrix and the one estimated for Eukaryotic phylogenetic analysis.

- a data set of 56 ribosomal proteins (7,112 sites *×* 86 taxa) described in Eme et al. (2023). To ensure computational tractability taxa were subsampled from the original 331 taxon dataset to maintain a representation of Asgard archaea, TACK archaea, and Euryarchaeota. A tree topology, denoted *T*_*R*_ estimated under LG+C60+G4, is used for comparing different exchangeabilities matrices.

We also used three empirical concatenated super-matrices to validate the empirical-estimated matrix discussed above to be used for Eukaryotic phylogenetic analysis and compare it with the LG matrix. For each data set, we consider two trees, the correct topology and an artifactual one (product of long-branch attraction artifacts). The data sets and trees are as follows:

- a data set of the 133-protein data set (24,291 sites *×* 40 taxa) described in (Brinkmann et al., 2005) to assess the placement of the Microsporidia in the tree of eukaryotes. The correct topology denoted *T*_*M*_, was originally recovered with LG+C20+F+G4, (Susko et al., 2018) and places the Microsporidia branch as sister to Fungi. The artifactual topology denoted *T*_*MA*_, was recovered with LG+F+G4 and groups the Microsporidia with the archaeal outgroup (i.e. branching sister to all other eukaryotes) due to an LBA artifact.
- a data set of 146 proteins (35,371 sites *×* 37 taxa) described in Lartillot et al. (2007) to assess the placement of the Nematodes in the animal tree of life. The correct topology, denoted *T*_*N*_, was recovered with LG+C20+F+G4 where Nematodes branch as sister to arthropods. The artifactual topology denoted *T*_*NA*_, was recovered with LG+F+G.
- a data set of 146 proteins (35,371 sites *×* 32 taxa) assembled in Lartillot et al. (2007) to assess the position of the Platyhelminths in the animal tree of life. The correct topology denoted *T*_*P*_, was recovered with CAT+GTR places Platyhelminths within the Protostomia. The artifactual topology, denoted *T*_*P A*_, recovered with LG+F+G4 and places Platyhelminths within Coelomata and many mixture models (see Lartillot et al. (2007); Susko et al. (2018); Wang et al. (2017))

## 4 Parameter Estimation Performance

To validate our implementation, we simulated 100 MSAs, with 10,000 sites and 10 taxa, using Alisim (Ly-Trong et al., 2021). Each alignment was simulated under the following conditions: LG exchangeabilities; a profile mixture model with 4 profiles, we arbitrarily chose the first 4 profiles from the C60 model, which turn out to be quite distinct (Fig. S2); a 10-taxon tree, depicted in Fig. S1 in the Supplement, obtained from the empirically estimated tree *T*_*M*_ defined above, after randomly removing taxa; and a discretized gamma distribution G4 (Yang, 1994), with *α* = 0.67, where *α* was chosen from an empirical data estimate (obtained after fitting the model LG+C60+G4 on the tree *T*_*M*_ for the Microsporidia data set). The arbitrarily chosen weights of the profiles were 0.35, 0.15, 0.25, and 0.25, respectively. For each MSA, we jointly estimated exchangeabilities, branch lengths, profile weights, and the rate parameter *α*. The only parameters not estimated were the tree topology and the profiles. We chose the POISSON exchangeability matrix, where all entries are equal to 1, as the initial values for the exchangeabilities to guarantee that the success of optimization was not due to the starting values being close to the true values.

Fig. 1(A) shows a histogram of the difference between true and estimated exchangeability deviation for all entries and for all 100 simulations. Particularly, this plot shows how most entries were accurately inferred since most of the mass is around zero. Fig. S3 in the supplement gives separate box plots for each exchangeability entry and shows that all entries are adequately estimated.

**Figure 1:**
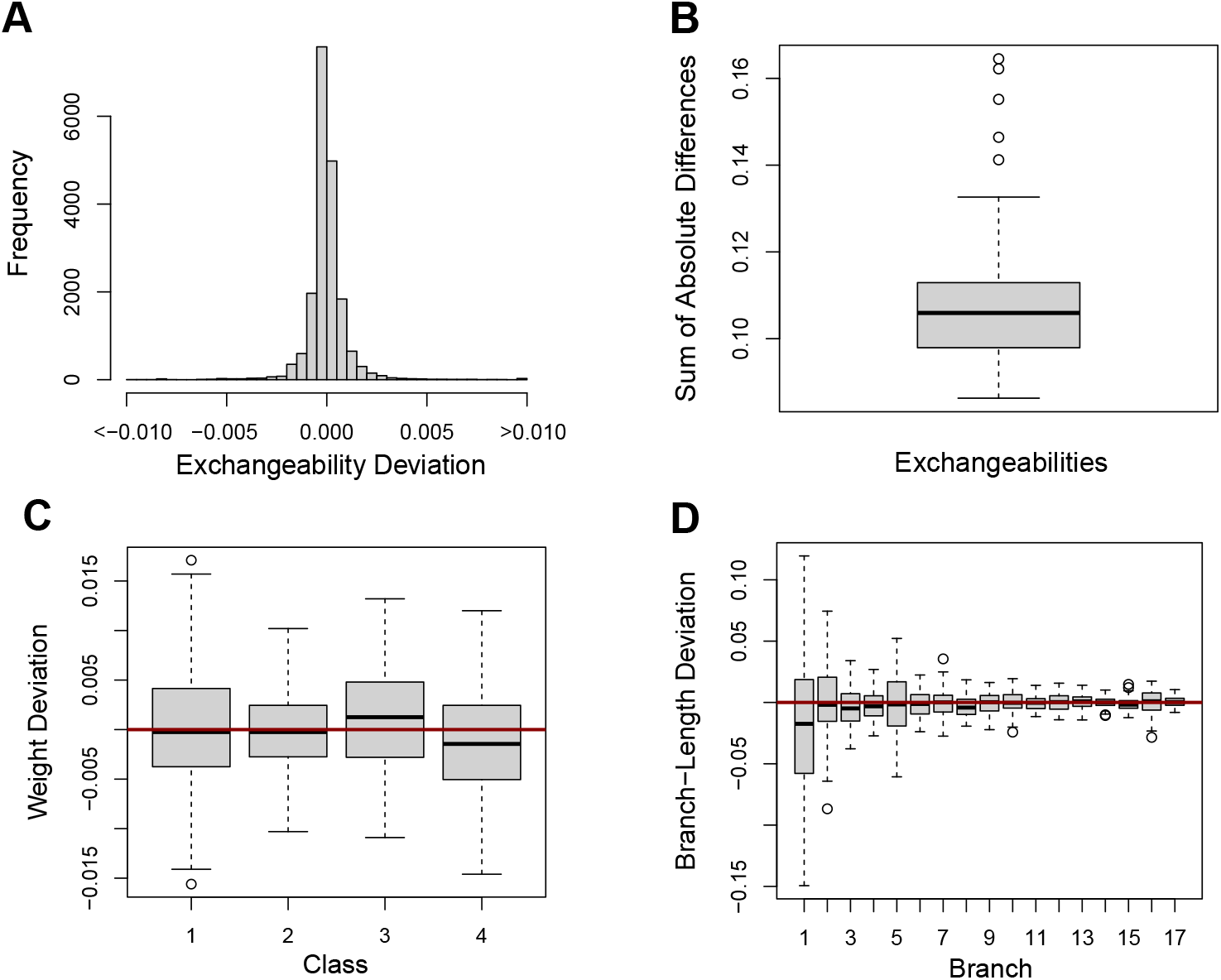
Plots showing the comparison between true and estimated parameters for all 100 simulations. For the box plots, a red line at zero represents a perfect estimation of the true parameters. (A) Histogram showing the differences between true and estimated exchangeability deviation for all entries. (B) Box plot showing the sum of absolute differences between the true (LG) and estimated exchangeabilities. (C) Box plot showing the differences between true and estimated weights for the four classes used to simulate the data. (D) Box plot showing the differences between true and estimated branch lengths. Branches are ordered from largest to shortest (the reason why variability decreases from left to right).

To investigate the performance of estimation of all exchangeabilities jointly, we compute for each estimated matrix *S*, the sum of absolute differences (SAD) between the true exchangeability matrix (LG) and *S*. Fig. 1(B) shows the box plot of the SAD for all simulations. For reference, the SAD between the true matrix (LG) and the starting matrix (POISSON) is *∼* 0.5, which is considerably larger than that of any estimated matrix. Moreover, the SAD between the LG matrix, and the mean estimated matrix is *∼* 0.02 (Fig. 2), suggesting consistency of the exchangeability estimation.

**Figure 2:**
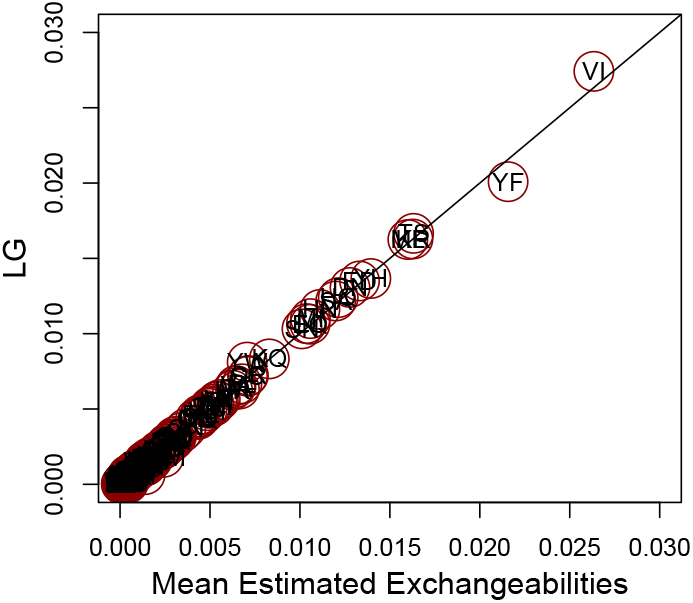
Entry-wise comparison between the mean exchangeabilities estimated from the simulated data and the LG exchangeabilities used to generate the data. Each dot represents an entry in the exchangeability matrix, where the *x*-coordinate is the mean estimated exchangeability over all 100 simulations and the *y*-coordinate is the corresponding LG entry. Each point is labeled with the two amino acids it represents.

Other parameters optimized jointly with exchangeabilities were also accurately estimated including profile weights (Fig. 1(C)), branch lengths (Fig. 1(D)), and the alpha shape parameters (Fig. S4).

Over 100 simulations, the median total CPU time used to estimate all parameters for each simulated dataset is 4297 seconds and the median wall-clock is 217 seconds on an Intel Xeon E5-2697 with 64GB RAM when using IQ-TREE2’s multi-threading option on 20 cores.

## 5 Estimating Exchangeabilities Improves Topological Accuracy

In (Baños et al., 2023), it was shown that misspecification of the exchangeabilities can severely hamper tree estimation. Specifically, it was shown that, under a ‘rich’ profile mixture model, data simulated under POISSON exchangeabilities and fitted using fixed LG exchangeabilities together with a profile mixture model that includes an ‘F-class’ (a profile that is defined from the empirical frequencies of amino acids from the MSA) performs poorly.

To determine if GTR estimation would address this problem, we investigated a similar scenario by simulating 50 MSAs of length 10,000 using the POISSON+C10+G4 {0.5} model on a 12-taxon tree shown in Fig. S5 (L). We separately fitted the profile mixture model C10+F+G4 {0.5} using the POISSON, LG, and GTR exchangeabilities, and two tree topologies, the correct tree and an artifactual one corresponding to the long-branch attraction (LBA) artifact (Fig. S5 (R)). For all models, branch lengths and weights of the profiles were estimated, and for GTR+C10+F+G4 {0.5} exchangeabilities were also estimated. Table 1 shows, for each model, the proportion of times the true tree had a higher log-likelihood than the LBA tree. As expected (Baños et al., 2023), LG+C10+F performs poorly when compared to the model used for simulation, POISSON+C10+F; GTR+C10+F, performs much better than LG and is much closer in performance to the model used for simulation, POISSON+C10+F. If more taxa and sites were considered, the GTR exchangeability estimates are expected to approach the true POISSON exchangeabilities and tree estimation would improve concomitantly.

**Table 1:**
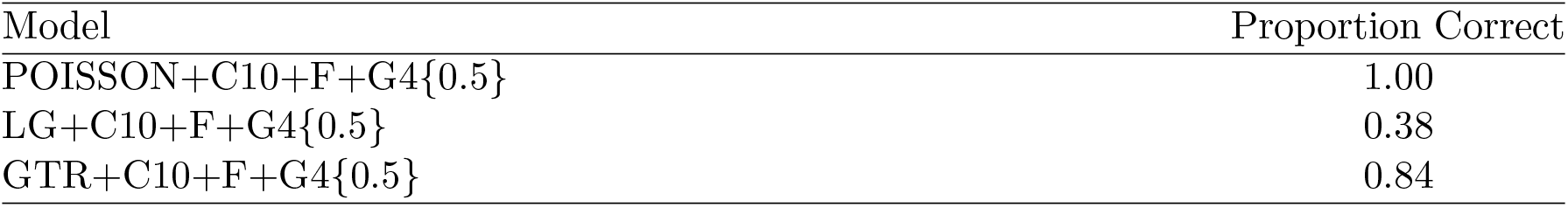
Proportion of times the correct tree is preferred over the artifactual LBA tree for 50 simulated MSAs simulated under the model POISSON+C10+G4*{*0.5*}*. The only difference between fitted models is the choice of exchangeabilities. For the model GTR+C10+F+G4*{*0.5*}* exchangeabilities are estimated using our implementation. McNe-mar’s test of the equality of proportions yielded a p-value *∼*0 when comparing the contin-gency table from trees for models GTR+C10+F+G4*{*0.5*}* and LG+C10+F+G4*{*0.5*}*.

| Model | Proportion Correct |
| --- | --- |
| POISSON+C10+F+G4{0.5} | 1.00 |
| LG+C10+F+G4{0.5} | 0.38 |
| GTR+C10+F+G4{0.5} | 0.84 |

## 6 Data analysis

We then investigated if GTRpmix improves model fit significantly compared to the LG matrix for the Microsporidia data set from Brinkmann and colleagues (Brinkmann et al., 2005). By fixing the tree topology *T*_*M*_ and the profiles of model C60, we jointly estimated the exchangeabilities of the GTR model, class weights, *α* from a discretized G4, and branch lengths. We refer to the estimated exchangeability matrix in this section as MXM. We compare this model against LG+C60+G4, where class weights, *α*, and branch lengths are estimated under fixed tree topology *T*_*M*_, profiles of model C60, and LG exchangeabilities. Table 2 shows the log-likelihoods, AIC, and BIC obtained from models LG+C60+G4 and MXM+C60+G4. Note that for MXM+C60+G4, 189 additional parameters are being estimated compared to LG+C60+G4. Nevertheless, the AIC and BIC scores suggest a preference for exchangeability estimation by a large margin (around 10k AIC and BIC units, Table 2).

**Table 2:**
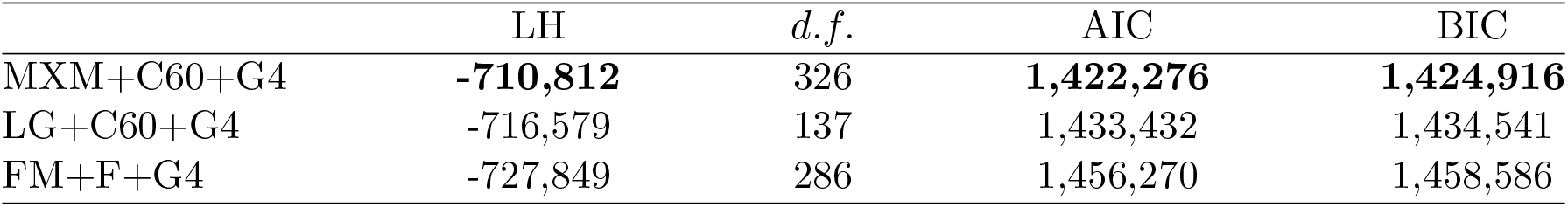
The log-likelihoods, degrees of freedom (denoted *d*.*f*., accounting for the 77 branch lengths in the tree), AIC, and BIC obtained from fitting models MXM+C60+G4, LG+C60+G4, and FM+F+G4. Both exchangeability matrices MXM and FM were estimated from the data, with the former estimated under C60 and the latter with the single frequency class ‘F’ estimated from the data.

Fig. 3 shows a comparison between the entries of the LG and the MXM matrix. Particularly, we see that many of the entries in the LG matrix are different than in the MXM matrix. To provide insight into these differences, we focus on a particular example. Some exchangeabilities involving cysteine (C) are increased in MXM compared to LG. This is likely because in C60 there is a profile (profile 8 as listed in IQ-TREE2) where cysteine has a frequency of *∼* 0.42 and both alanine and serine each have a frequency of *∼*0.18; these three amino acids account for almost 80% of the overall amino acid proportion of this profile. By excluding from C60 this and profile 4 (where cysteine has a frequency of 0.16 and only A, L, S, T, and V have a frequency greater than 0.05), the mean frequency of cysteine in the 58 remaining profiles is 0.009. Assuming this profile mixture is closer to reality, even if the exchangeabilities involving cysteine are non-negligible, a frequency profile with a substantial probability of cysteine is unlikely.

**Figure 3:**
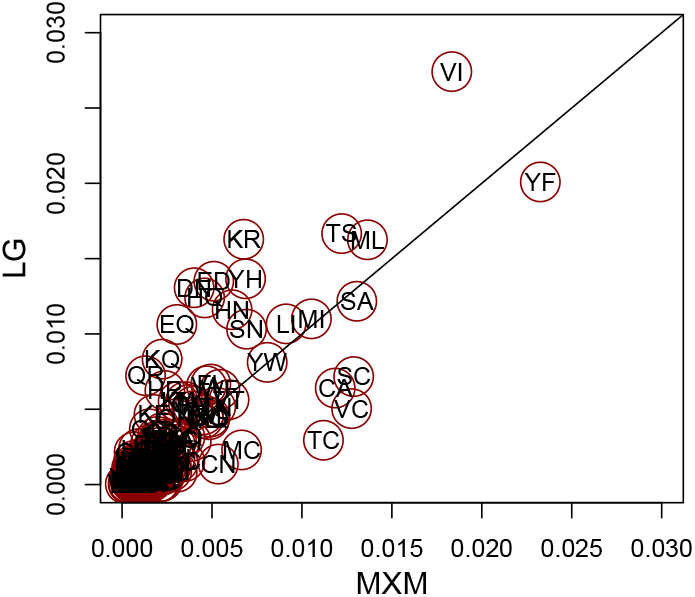
Entry-wise comparison between the MXM matrix, obtained from fitting a GTR matrix to the Microsporidia data set, and the LG exchangeabilities. Each dot represents an entry in the exchangeability matrix, where the *x*-coordinate is the entry of the MXM matrix and the *y*-coordinate is its corresponding LG matrix entry. Each point is labeled with the two amino acids it represents.

Thus a small proportion of sites with a cysteine and some other amino acid is expected. Because MXM takes frequency profiles into account it can recognize this. However, when a single profile is assumed in fitting, as with LG, a small exchangeability provides the only way to account for a low frequency of sites with cysteine and some other amino acids. A bubble plot of the differences between the LG and MXM exchangeabilities can be found in Fig. S6 in the supplement.

For comparison, we also estimated a GTR matrix, denoted FM, under a single profile and discretized G4 rates. Specifically, for the profile we used the overall frequencies of amino acids in the data set, also referred to as the ‘F-class.’ Fig. S7(L) depicts a comparison between the FM and MXM matrices. This figure shows, as expected, how MXM has higher cysteine exchangeabilities than FM, as was noted in the MXM to LG comparison. For additional comparison, the SAD between the FM and LG is *∼* 0.15, while the SAD between the FM and MXM matrices is *∼* 0.26. This means that the FM matrix is more alike to the LG matrix than the MXM matrix. Table 2 shows that the AIC, and BIC scores of both LG and FM when combined with C60 are worse than MXM underscoring the importance of fitting multiple profiles and exchangeabilities jointly.

The total running time for full estimation of the exchangeabilities for the MXM model, branch lengths, C60 class weights, and the shape parameter, *α* was *∼* 11 hours on 40 cores of an AMD EPYC 7543 Processor with 2T of RAM. We investigated whether accurate estimation of exchangeabilities would be possible by fixing branch lengths and the shape parameter *α* from G4. Specifically, we estimated the exchangeability matrix, denoted MXM^*F ix*^, by fixing branch lengths and *α* to the averages of these parameters obtained from fitting POISSON+C60+G4 and LG+C60+G4. The total running time to estimate MXM^*F ix*^ was *∼* 7 hours on the same computer and the resulting SAD between MXM and MXM^*F ix*^ is 0.017, suggesting the estimates are very similar (Fig. S7(R) in the supplement shows an entry-wise comparison between these two matrices). The log-likelihood obtained after fitting the model MXM^*F ix*^+C60+G4 (where branch lengths and *α* are re-estimated) is only 15 likelihood units greater than the one obtained from fitting MXM+C60+G4 with all parameters estimated simultaneously. This suggests that fixing branch lengths and *α* does not greatly affect the estimation of exchangeabilities while yielding important computation time savings.

## 7 Empirically Estimated Exchangeability Matrices

Since the estimation of exchangeabilities is computationally expensive even after fixing branch lengths and rates, many users will not have the computational resources to optimize matrices for their datasets of interest. Alternatively, users may have datasets that are not large enough to permit accurate exchangeability estimation (e.g., a single-protein alignment). For these reasons, we have estimated two exchangeability matrices from large data sets using the C60+G4 model that can be used as fixed matrices for phylogenetic analyses under profile mixture models. The first matrix we introduce is tailored for phylogenetic analyses of proteins encoded by eukaryotic nuclear genes and the other is for reconstructing relationships between nuclear-encoded proteins in eukaryotes and orthologs in Archaea.

### 7.1 The ELM Exchangeability Matrix for Eukaryotic Analyses

We estimated an exchangeability matrix, which we refer to as the **E**ukaryotic **L**inked **M**ixture (ELM) matrix. This matrix was estimated from the 78-taxon Pan-Eukaryotic data set described in the section ‘*Data Sets*’ above. We used the profiles from model C60, discretized G4 rates, and a tree topology recovered by fitting LG+C60+G4 to the data depicted in Fig. S10 in the supplement. To reduce computational time we used the approach described for MXM^*fix*^ estimation above; i.e., we fixed branch lengths and *α* to their averages based on estimates from LG+C60+G4 and POISSON+C60+G4. Thus for the ELM matrix estimation, we only optimized exchangeabilities and C60 profile weights jointly.

Fig. S8 in the supplement shows a bubble plot of the difference between the LG and ELM exchangeabilities, and Fig. S9(L) depicts an analogue of Fig. 3 for these two matrices. To compare the fit between these two matrices, we used the three empirical data sets (Microsporidia, Nematode, and Platyhelminths) described in the subsection ‘*Data Sets for Comparisons*’. Figure 4 contains, among other things, the likelihoods obtained from fitting models LG+C60+G4 and ELM+C60+G4 for the correct and artifactual topologies of each data set. Note that, by applying the KHns test developed in (Susko, 2014) to compare two fixed tree topologies, a likelihood difference between fitting the true and artifactual tree is considered significant, with a 5% significance level, if it is greater than 5.53 for the Microsporidia, 2.99 for the Nematode, and 1.92 for the Platyhelminths data sets. Clearly, the ELM matrix produces considerably better likelihood scores for all data sets than the LG matrix. We note that for the Platyhelminths data set the ELM+C60+G4 significantly prefers the true tree over the artefactual tree, whereas the LG+C60+G4 matrix does not. For the other two datasets, all models preferred the true tree, although the LG model consistently had the weakest preference, Figure 4.

**Figure 4:**
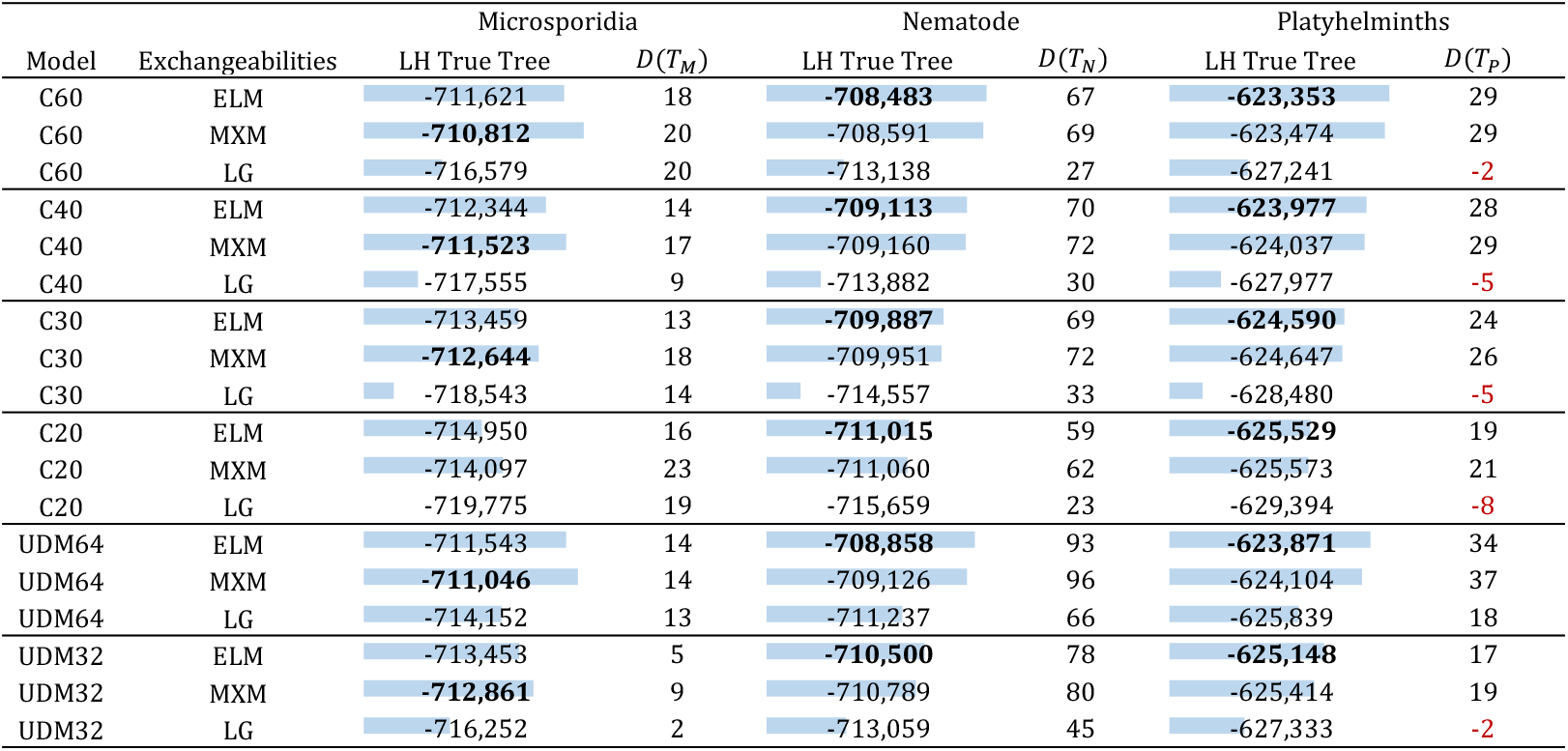
The log-likelihoods of the trees estimated from the three empirical data sets and the difference between fitting the true and the artifactual tree for each data set. The bar scale goes from the lower value per column (empty) to the highest value (full) per column. *D*(*T*_*X*_) denotes the log-likelihood of the ‘correct tree’ (e.g. *T*_*M*_) minus the ‘incorrect’ tree (e.g. *T*_*MA*_). Positive values of *D*(*T*_*X*_) reflect a preference for the true tree, while negative (shown in red) preference for the LBA tree. The best LH score per model and data set is shown in bold.

We also found that even when the ELM matrix was optimized using the profiles from C60, it still provided better fit and improved topological accuracy when fitting different sets of profiles. Figure 4 contains the likelihoods for the three data sets when fitting the profiles in C40, C30, C20, UDM32, and UDM64 with LG and ELM exchangeabilities. For all profile mixture models, the ELM matrix provides better likelihood scores than the LG matrix. We also see that the ELM matrix always prefers the true tree, which is not true for the LG matrix under models C20, C40, and UDM32 for the Platyhelminths data set.

To make a broader comparison, we also looked at the likelihoods for all the profile mixture models used in the comparisons above with MXM exchangeabilities, shown in Figure 4. As expected, independently of the profiles, for the Microsporidia data set the MXM matrix produces better likelihood scores since this matrix was optimized on this data set. Nonetheless, for the other two data sets the ELM matrix produces better likelihood scores.

To confirm that the choice of branch lengths and *α* did not negatively affect the estimation of the ELM matrix, we also estimated an exchangeability matrix with the C60+G4 model, referred to as ELM50, on the 50-taxon Pan-Eukaryotic matrix described in the ‘*Data Sets*’ section. We chose this smaller data set to ease some of the computational burden so that joint exchangeability, profile weight, branch length, and *α* estimation could be conducted. The entry-wise comparison between the ELM and ELM50 matrices in Fig. S9(R) shows that these two matrices are in general very similar (the SAD between these matrices is 0.0.34). Table S1 in the supplement shows the equivalent of Figure 4 for the ELM50 matrix. We note that in all cases the ELM matrix produces better likelihood scores than the ELM50 matrix. We conclude that estimating exchangeabilities using fixed branch lengths and *α* did not affect negatively the estimation of the ELM matrix. Using fixed parameters may allow many more taxa to be used which should improve estimation. However, in cases where the guide tree and parameters, are far from optimal, it could lead to poor estimation.

On the other hand, the performance of the ELM matrix under a single profile model is subpar compared to the LG matrix (Figure 5). With a single ‘F-class’ for each data set, both ELM and MXM perform worse than LG, so it is inadvisable to use these models as part of a site-homogeneous model with a single profile. Although LG performs better than the other matrices for this site-homogeneous setting, it does considerably worse than these matrices when used with any of the mixture models in Figure 4.

**Figure 5:**
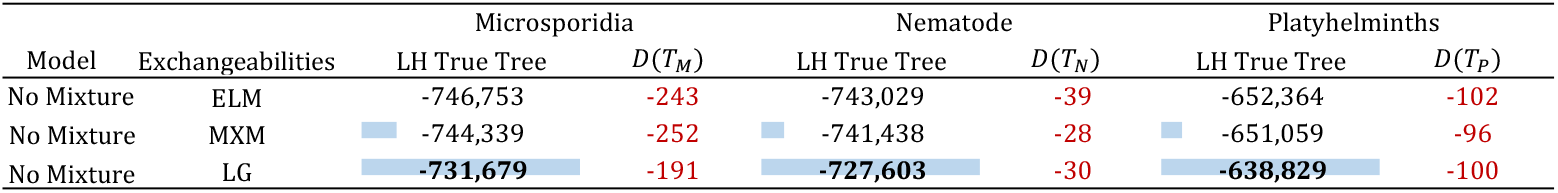
Similar to the table in Fig. 4 but for models with a single profile, denoted as F, estimated from the overall frequencies of amino acids in each data set.

### 7.2 The EAL Exchangeability Matrix for reconstructing relationships between eukaryotes and Archaea

We estimated an exchangeability matrix, which we refer to as the **E**ukaryotic and **A**rcheal **L**inked mixture (EAL) matrix, from the 86-taxon Eukaryotic-Archaeal data set described in the section ‘*Data Sets*’ above. We used the profiles from model C60, discretized G4 rates, and the tree topology depicted in Fig. S10.5 in the supplement, which is assumed to be the correct one. Similar to how we estimated the matrix MXM^*fix*^, we fixed branch lengths and *α* to their average from fitting models LG+C60+G4 and POISSON+C60+G4.

Fig. S11 and Fig. S12 in the supplementary material show a comparison of the EAL, ELM, and LG matrices. These plots indicate that these matrices are all quite different, with the EAL matrix being somewhat more similar to ELM than to LG. The SAD from ELM and EAL is *∼* 0.19, whereas the SAD between LG and EAL is *∼* 0.26. To show this matrix gives a better fit for data sets with both eukaryotic and archaeal sequences, we used the 56 ribosomal protein data set and the tree *T*_*R*_ described in the *Data sets* section. Table 3 shows the log-likelihood obtained by fitting models LG+C60+G4, ELM+C60+G4, and EAL+C60+G4 for this data set and tree. It is clear that EAL produces the best likelihood score by a wide margin. Note that since all the models have the same number of parameters, their fit can be directly compared using the log-likelihood scores (under these conditions, AIC and BIC will yield identical orderings of relative model fit to log-likelihood comparisons).

**Table 3:**
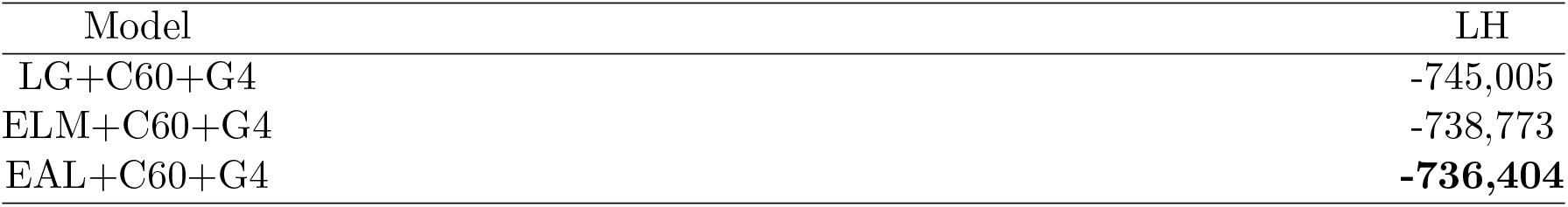
The log-likelihoods obtained from fitting models LG+C60+G4, ELM+C60+G4, and EAL+C60+G4 on the for the 56 ribosomal protein data set and the tree *T*_*R*_.

| Model | LH |
| --- | --- |
| LG+C60+G4 | -745,005 |
| ELM+C60+G4 | -738,773 |
| EAL+C60+G4 | **-736,404** |

## 8 Conclusion

We have shown that estimation of linked exchangeabilities jointly with profile mixture model weights in the GTRpmix model framework provides substantially better model fit for empirical amino acid alignments, and it can improve topological estimation for especially difficult problems where widely used empirical exchangeability matrices like LG fail. The GTRpmix model will be extremely useful for researchers investigating deep phylogenetic problems where the use of well-fitting site-heterogeneous models is especially important to avoid phylogenetic artifacts. Furthermore, we provide a number of pre-estimated matrices for use with profile mixture models in the analysis of eukaryotic nucleus-encoded protein data sets (e.g. MXM, ELM) and eukaryote-archaeal data sets (e.g. EAL).

Matrices ELM and EAL are available to use in the IQ-TREE2 software version v2.3.1. These matrices, together with MXM and ELM50, are also available in the Github repository https://github.com/RogerLab/GTRPMIX/. Additionally, in the supplement we have included an IQ-TREE2 sample command line to estimate exchangeabilities as it was done in this work.

## 9 Acknowledgments

We thank Dominik Schrempf for insightful discussions and advice. This work, J.D., and H.B. were supported by the Moore-Simons Project on the Origin of the Eukaryotic Cell, Simons Foundation grant 735923LPI (DOI: https://doi.org/10.46714/735923LPI) and by NSERC Discovery Grants awarded to A.J.R. and E.S. H.B. was also partially supported by NSF grant DMS 2331660. This work was supported in part by the National Science Foundation (NSF) Division of Environmental Biology (DEB) through the grant 2100888 awarded to M.W.B. L.E. was also supported by the European Research Council (ERC Starting Grant No 803151). Chan-Zuckerberg Initiative grant for open-source software for science [to B.Q.M. and R.L.]; and an Australian Research Council Discovery Grant [DP200103151 to R.L. and B.Q.M]. Additionally, H.B. would like to thank Balagopal Pillai for all the technical support.

