## Supplementary Material for "GTRpmix: A linked general-time reversible model for profile mixture models"

(2024), **0**, 0, *pp.* 1–**??**
/"Supplementary material˙GTR"

### Supplementary Material: Estimating Linked Exchangeabilities for Profile Mixture Models

Hector Banos*[,1,2], Thomas KF. Wong[3], Justin Daneau[2], Edward Susko[2], Bui Quang Minh[4], Robert Lanfear[5], Matthew W. Brown[6], Laura Eme[7], and Andrew J. Roger[2]

[1] *Department of Mathematics and Statistics, Faculty of Science, Dalhousie University, Halifax, NS, Canada.*

[2] *Department of Biochemistry and Molecular Biology, Faculty of Medicine, Dalhousie University, Halifax, NS, Canada.*

[3] *Research School of Biology, Australian National University, Australian Capital Territory, Australia.*

[4] *School of Computing, College of Engineering and Computer Science, Australian National University, Canberra, ACT 2600, Australia.*

[5] *Ecology and Evolution, Research School of Biology, College of Science, Australian National University, Canberra, ACT 2600, Australia.*

[6] *Department of Biological Sciences, Mississippi State University, Mississippi State, MS, USA.*

[7] *Université Paris-Saclay, Laboratoire d'Ecologie, systématique et Evolution, Gif-sur-Yvette, France*

#### *Data Set*

To estimate the "Eukaryote" exchangeability matrix we selected a 240-protein data et with 76,840 sites and 78 taxa as a taxonomically representative subsample of all eukaryotes in the PhyloFisher database Tice et al. (2021). As mentioned in the manuscript, taxa were selected based on their known membership in a particular higher-level eukaryotic taxon and their phylogenetic position. Additionally, we included in this matrix *Microheliella maris* an orphan protistan taxon recently shown to occupy an important phylogenetic position amongst the Diaphoretickes Yazaki et al. (2022). Data from this taxon was not available or included in the original PhyloFisher database and thus orthologs were collected from *M. maris* using the PhyloFisher algorithm.

As also mentioned in the manuscript, we also looked at a smaller subset of the PhyloFisher database consisting of a 240-protein data set with 77,965 sites and 50 taxa. The 50 taxon data only differs from the 78 taxa data set by the exclusion of taxa within

© The Author 2024. Published by Oxford University Press. All rights reserved. For permissions, please each sampled taxonomic suprakingdom level group Burki et al. (2020).

As mentioned in the manuscript, to estimate the exchangeability matrix for reconstructing relationships between eukaryotes and Archaea, we used a 54-protein data set with 14,704 sites and 86 taxa. This data set includes a subset of the taxa presented in Eme et al. (2023) Eme et al. (2023). Taxa were sampled with a selection of eukaryotes, Asgard archaea, TACK archaea, and Euryarchaeota. Treemmer Menardo et al. (2018) was used for the taxon sampling, followed by manual curation taking into account proteome completeness.

To compare the fit of the new empirically-estimated matrices to be used for reconstructing relationships between eukaryotes and Archaea against the LG matrix and the one estimated for Eukaryotic phylogenetic analysis, we used a data set of 56 ribosomal proteins (7,112 sites $\times$ 86 taxa) described in Eme et al. (2023). For this data set, we first discarded DPANN archaea, since they are the most distantly related to eukaryotes and tend to present long branches and compositional biases due to their probable symbiotic lifestyle Dombrowski et al. (2019). We then subsampled 86 representatives among the remaining taxa using Treemmer v.0.3 Menardo et al. (2018).

#### *Supplementary Figures, Tables, and Sections*

All the materials are presented in the order they are referred to in the manuscript.

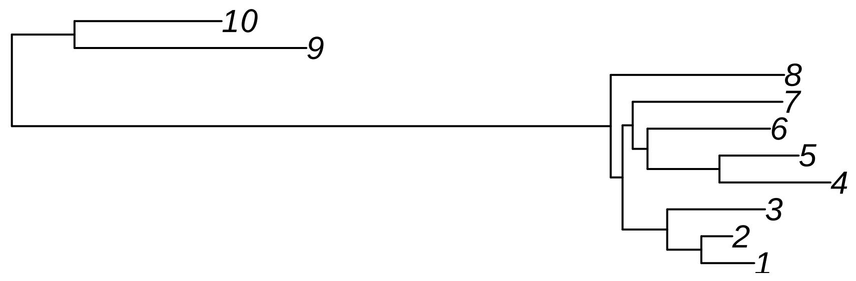


Fig. S1. The 10-taxon tree used to simulate data for the section '*Parameter Estimation Performance.*' This tree was obtained after randomly removing taxa form the empirically estimated tree $T_M$ until 10 taxa remained. Specifically such tree is:
(((((1:0.127073,2:0.074368):0.08191,3:0.234948):0.107157,(((4:0.266625,5:0.190106):0.172775,6:0.294129):0.035602,
7:0.35961):0.024291):0.028459,8:0.416453):1.437843,(9:0.556753,10:0.353276):0.150375);

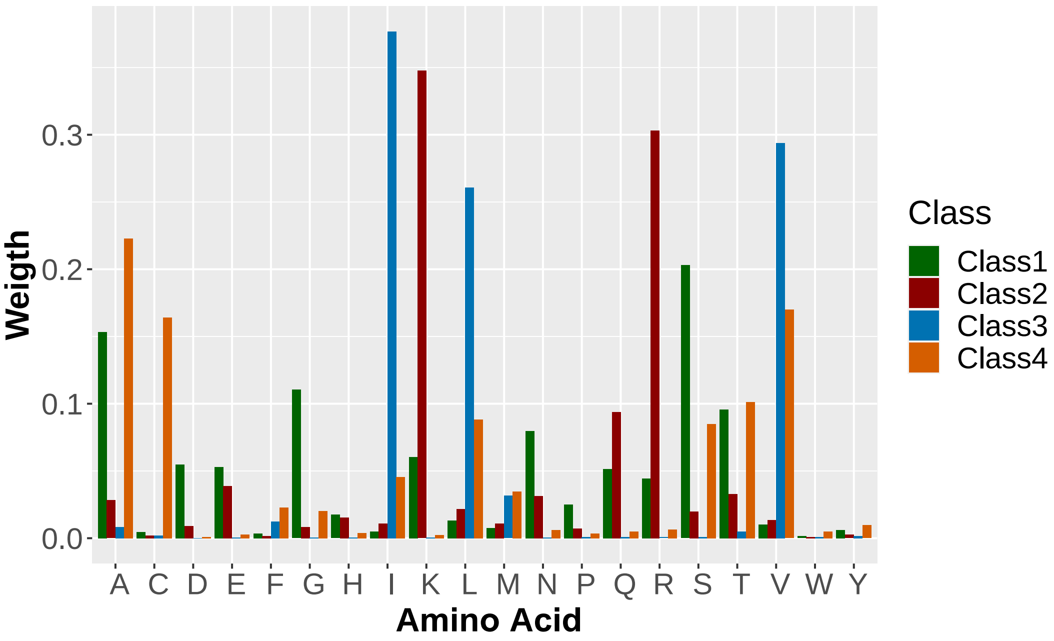


Fig. S2. The four amino acid frequency vectors (profiles) of the model used to simulated the data for the section '*Parameter Estimation Performance.*' This classes the first four profiles from the model C60.

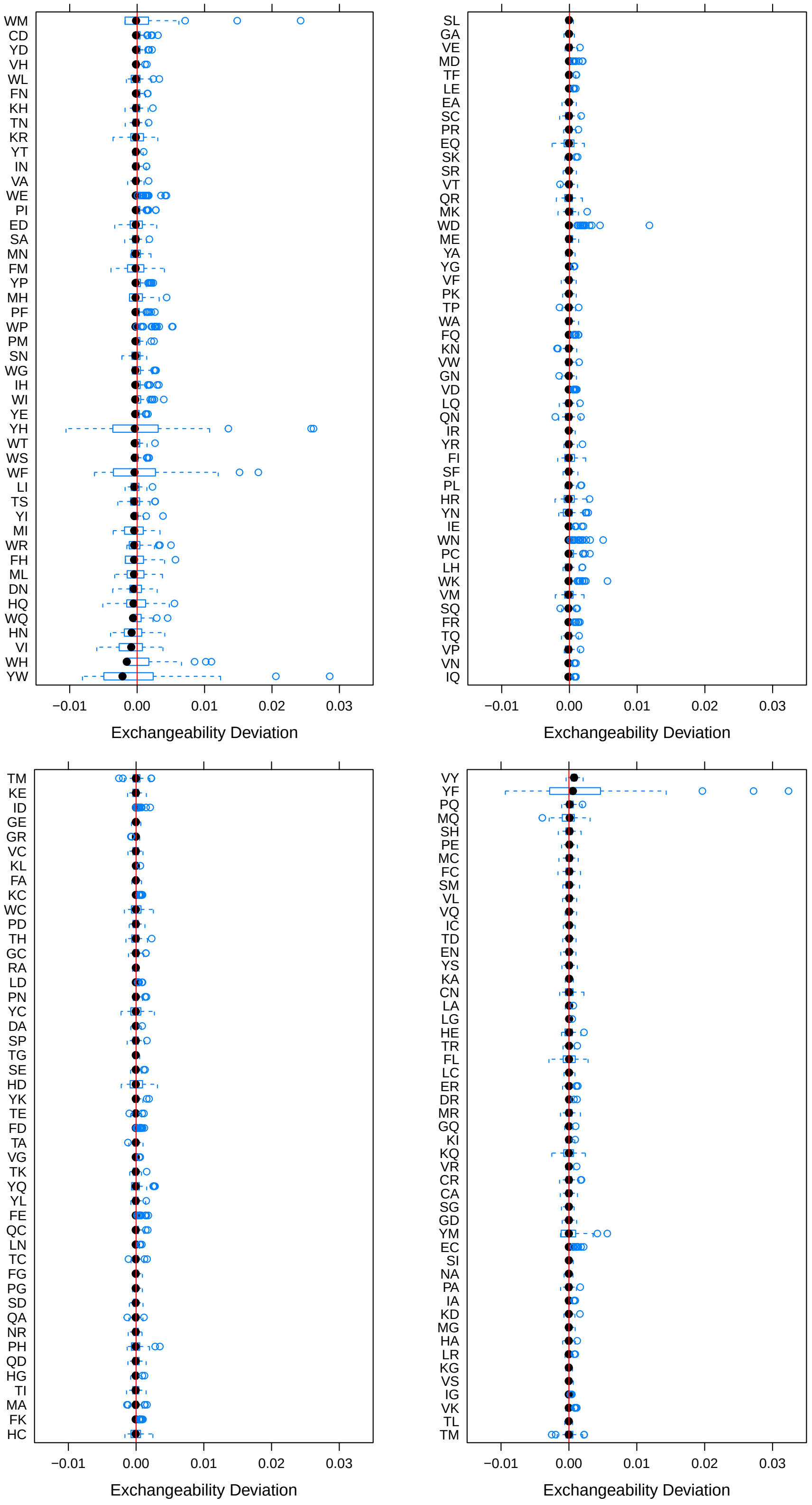

Fig. S3. Box plots showing the differences between true and estimated exchangeabilities for all 190 entries. Exchangeabilities are divided in 4 plots for better readability. The mean difference between the true and estimated $\alpha$ is $\sim$0.004 with a variance of $\sim$0.0003.

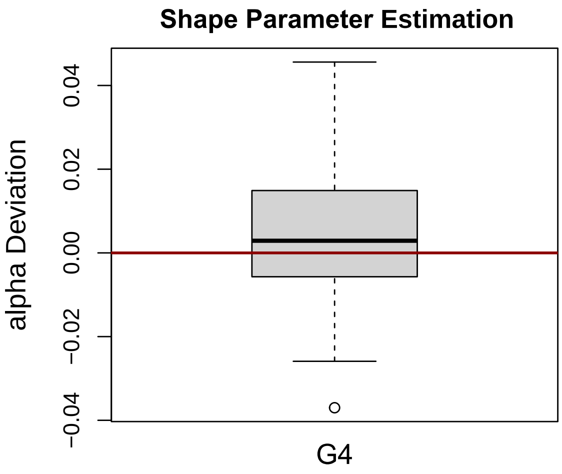


Fig. S4. Box-plot showing the differences between true and estimated $\alpha$.

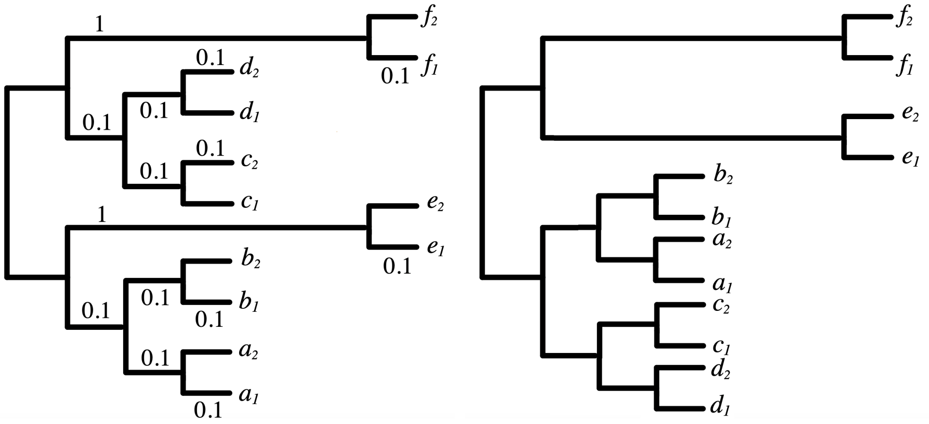


Fig. S5. (Left)The tree used for the simulations described in the section *'Improving Topological Accuracy.'* This tree is known to be susceptible to LBA artifacts under model misspecification. (Right). The artifactual tree topology corresponding to the long-branch attraction (LBA) biases obtained from data generated frp, the tree on the left.

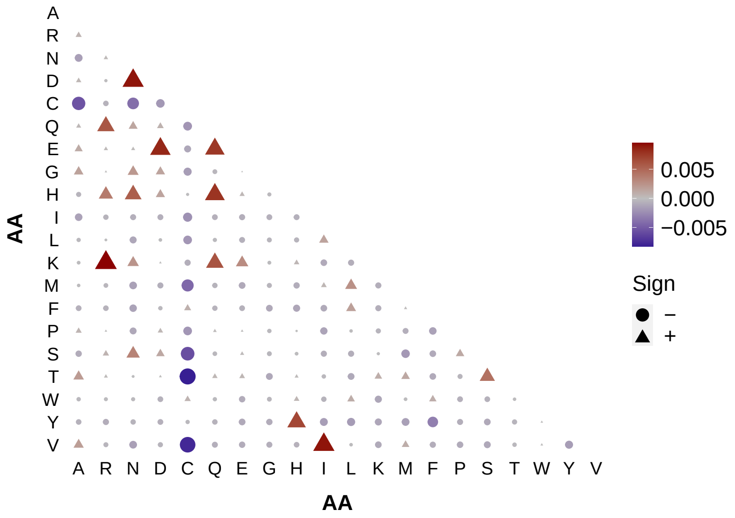


Fig. S6. A bubble/heat plot showing the difference between the lower diagonal of the LG matrix minus that of the MXM matrix. Small figurines represent both matrices having similar entries. Red-ish triangles represent the LG having higher entries than MXM while blue-ish circles represent the exact opposite.

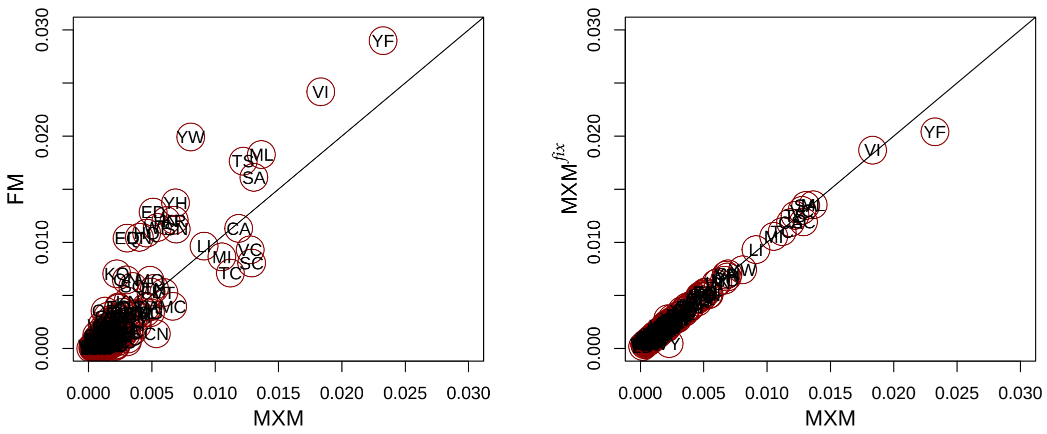


Fig. S7. (L) Entry-wise comparison between the MXM exchangeabilities estimated from the miscrosporidia data set via the GTR+C60+G4 model and the FM exchangeabilities estimated from the same data via the GTR+F+G4 model. Each dot represents an entry in the exchangeability matrix, where the $x$-coordinate is and entry of MXM and the $y$-coordinate is the corresponding FM entry. Each point is labeled with the two amino acids it represents. (R) A similar plot as the one in the left but comparing matrices MXM and MXM$^{fix}$ estimated after fixing branches and $\alpha$ as describe in the main manuscript.

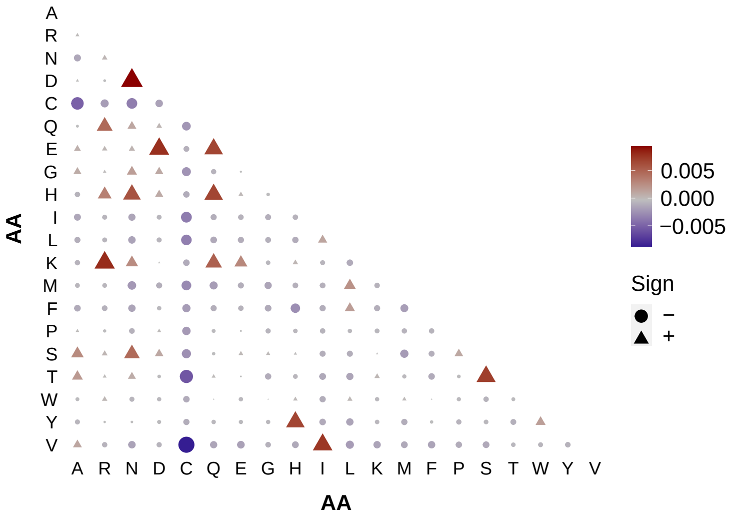


Fig. S8. A bubble/heat plot showing the difference between the lower diagonal of the LG matrix minus that of the ELM matrix. Small figurines represent both matrices having similar entries. Red-ish triangles represent the LG having higher entries than ELM while blue-ish circles represent the exact opposite.

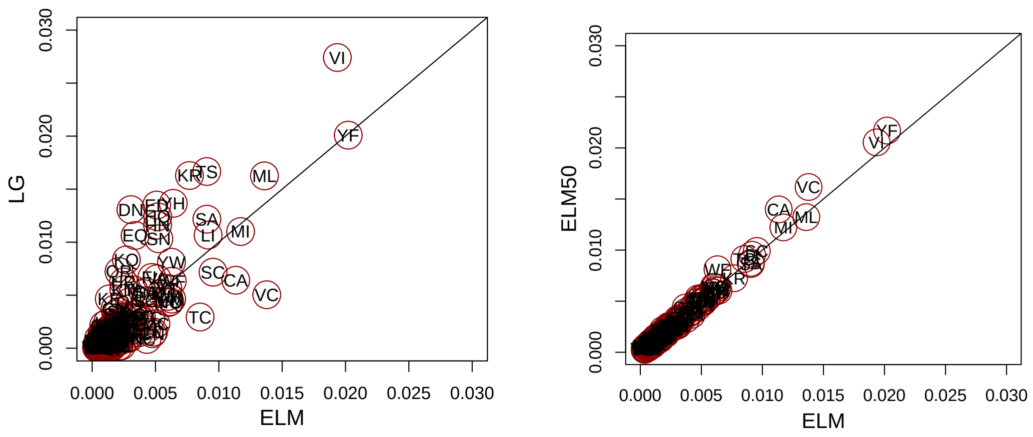


Fig. S9. (L) Similar to the plots in Figure S6 but comparing the ELM, estimated from the 78-taxon matrix described in the section '*Data Sets*' via the GTR+C60+G4 model, and the LG exchangeabilities. (R) A similar plot as the one in the left but comparing matrices ELM and ELM50, estimated from the 50-taxon matrix described in the section '*Data Sets*' via the GTR+C60+G4 model after fixing branch lengths and $\alpha$ as described in the manuscript.

| | | Microsporidia | | Nematode | | Platyhelminths | |
|---|---|---|---|---|---|---|---|
| Model | Exchangeabilities | $L(T_M)$ | $D(T_M)$ | $L(T_N)$ | $D(T_N)$ | $L(T_P)$ | $D(T_P)$ |
| C60 | ELM50 | -711,688 | 19 | -708,563 | 66 | -623,423 | 28 |
| C40 | | -712,385 | 15 | -709,181 | 69 | -624,036 | 27 |
| C30 | | -713,513 | 13 | -709,960 | 68 | -624,654 | 23 |
| C20 | | -714,992 | 18 | -711,114 | 58 | -625,618 | 18 |
| UDM64 | | -711,509 | 14 | -708,903 | 92 | -623,905 | 33 |
| UDM32 | | -713,403 | 6 | -710,524 | 77 | -625,168 | 16 |
| No Mixture | | -745,979 | -240 | -742,435 | -35 | -651,881 | -100 |

Table S1. The log-likelihoods of the trees estimated from the three empirical data sets and the difference between fitting the true and the artifactual tree for each data set. $D(T_X)$ denotes the log-likelihood of the 'correct tree' (e.g. $T_M$) minus the 'incorrect' tree (e.g. $T_{MA}$).

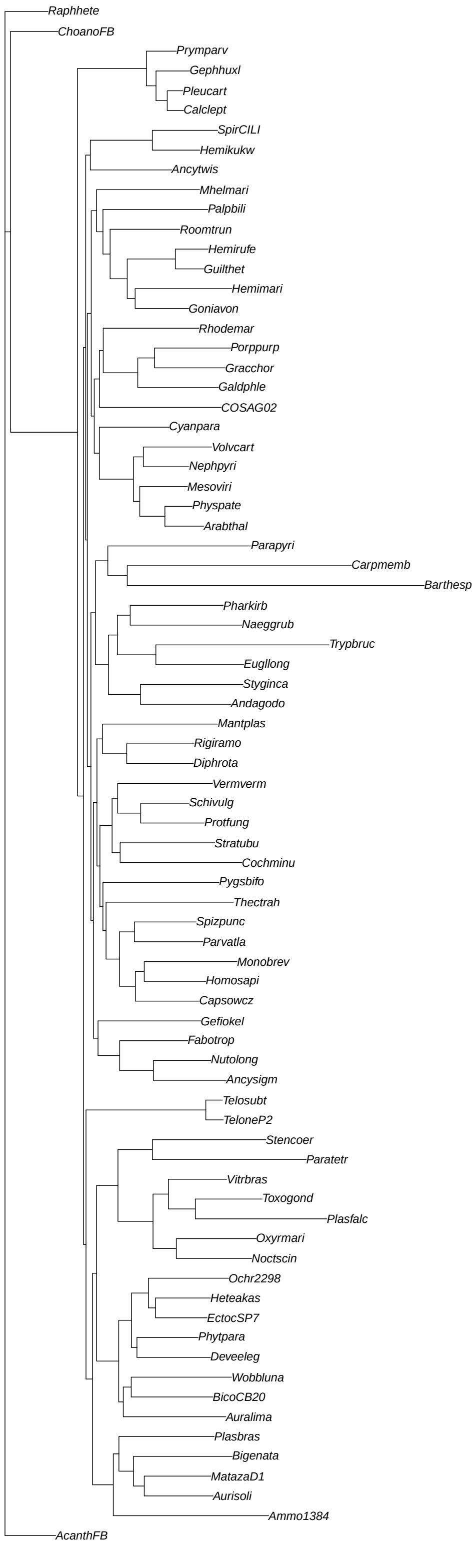


Fig. S10. The unrooted tree topology used to estimate the ELM matrix.

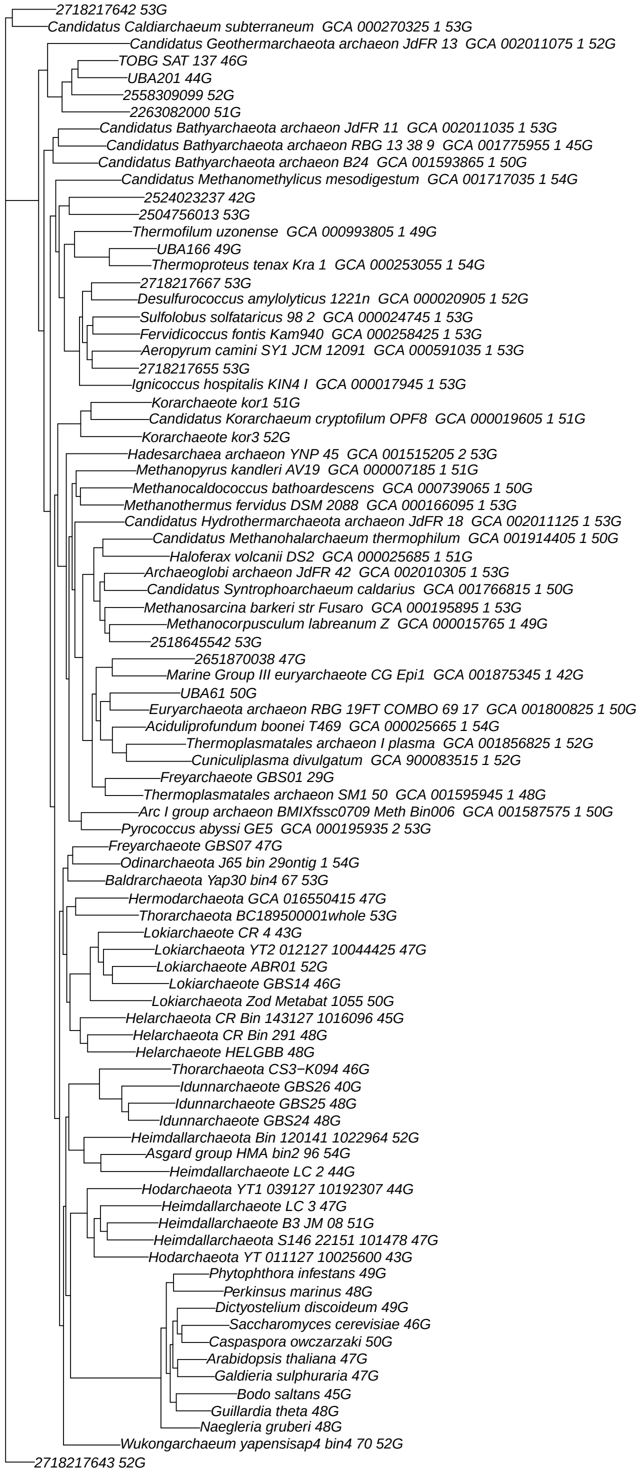


Fig. S10.5. The unrooted tree topology used to estimate the EAL matrix.

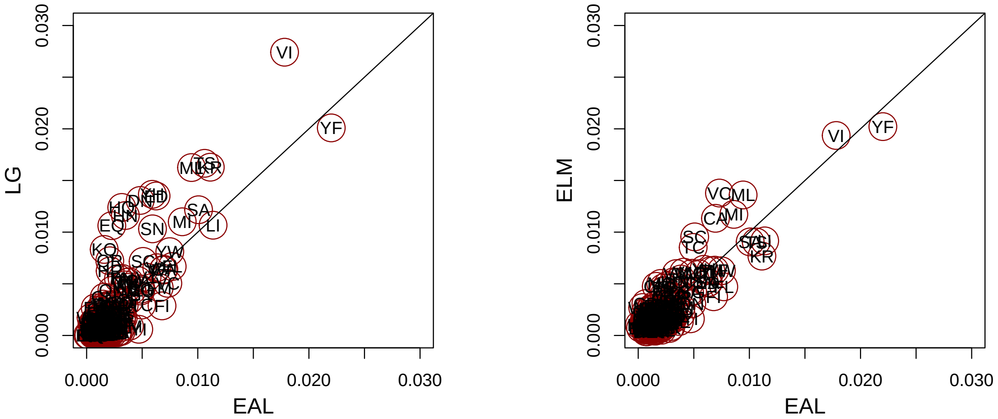


Fig. S11. (L) Entry-wise comparison between the EAL exchangeabilities and the LG exchangeabilities. Each dot represents an entry in the exchangeability matrix, where the $x$-coordinate is and entry of EAL and the $y$-coordinate is the corresponding LG entry. Each point is labeled with the two amino acids it represents. (R) A similar plot as the one in the left but comparing matrices EAL and ELM.

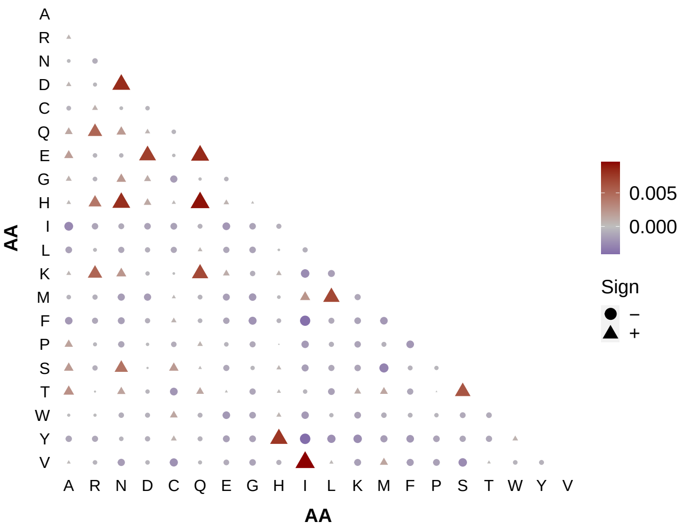


Fig. S12. A bubble/heat plot showing the difference between the lower diagonal of the LG matrix minus that of the EAL matrix. Small figurines represent both matrices having similar entries. Red-ish triangles represent the LG having higher entries than EAL while blue-ish circles represent the exact opposite.

#### *Sample command line*

We present an example of the command line used to estimate exchangeabilities for a profile mixture model 'C60+G4' but any profile mixture model and rates can be used. To estimate a single set of linked exchangeabilities, in the model definition the matrix

'GTR20' must be specified (resp. GTR for nucleotide data) together with the flag '–link-exchange-rates.' While a guide tree is not needed, we highly recommend using a fixed tree topology to estimate exchangeabilities. Since matrix estimation can be time-consuming, we also recommend using the flag '-me 0.99' to reduce the optimization threshold for faster optimization.

```
iqtree -s <alignment> -m GTR20+C60+G4 --link-exchange-rates -te
<guide_tree> -me 0.99 -mwopt
```

The user can determine the starting exchangeabilities before optimization. Choosing adequate exchangeabilities can make estimation considerably faster. For example:

```
iqtree -s example.phy -m GTR20+C60+G4 --link-exchange-rates --gtr20-model
LG -te <guide_tree> -me 0.99 -mwopt
```

specifies the LG matrix as the starting matrix via the flag '–gtr20-model' (the default starting matrix is POISSON, i.e. equal exchangeabilities). For this flag, the user can specify any matrix, even those matrices defined by the user via the '-mdef' flag. If the user is agnostic of the exchangeabilities, we recommend using the default matrix (although it can be time-consuming).
